# DNA Methyltransferase Inhibitors with Novel Chemical Scaffolds

**DOI:** 10.1101/2020.10.13.337709

**Authors:** K. Eurídice Juárez-Mercado, Fernando D. Prieto-Martínez, Norberto Sánchez-Cruz, Andrea Peña-Castillo, Diego Prada-Gracia, José L. Medina-Franco

## Abstract

Inhibitors of DNA methyltransferases (DNMTs) are attractive compounds for epigenetic drug discovery. They are also chemical tools to understand the biochemistry of epigenetic processes. Herein, we report five distinct inhibitors of DNMT1 characterized in enzymatic inhibition assays that did not show activity with DNMT3B. It was concluded that the dietary component theaflavin is an inhibitor of DNMT1. Two additional novel inhibitors of DNMT1 are the approved drugs glyburide and panobinostat. The DNMT1 enzymatic inhibitory activity of panobinostat, a known pan inhibitor of histone deacetylases, agrees with experimental reports of its ability to reduce DNMT1 activity in liver cancer cell lines. Molecular docking of the active compounds with DNMT1, and re-scoring with the recently developed Extended Connectivity Interaction Features approach, had an excellent agreement between the experimental IC_50_ values and docking scores.

## 1. Introduction

Historically, the term “epigenetics” is rooted in Waddington and Nanney’s work, where it was initially defined to denote a cellular memory, persistent homeostasis in the absence of an original perturbation or an effect on cell fate not attributable to changes in DNA.^[1–2]^ However, “epigenetics” is now used with multiple meanings, for instance, to describe the heritable phenotype (cellular memory) without modification of DNA sequences,^[3]^ or the mechanism in which the environment conveys its influence to the cell, tissue or organism.^[4]^ Regardless of the different definitions, the interest in epigenetic drug discovery increases, as revealed by the multiple approved epigenetic drugs or compounds in clinical development for epigenetic targets.^[5–6]^

DNA methyltransferases (DNMTs) are one of the primary epigenetic modifiers. This enzyme family is responsible for promoting the covalent addition of a methyl group from *S*-adenosyl-L-methionine (SAM) to the 5-carbon of cytosine, mainly within CpG dinucleotides, yielding *S*-adenosyl-*L*-homocysteine (SAH).^[7]^ DNMT1, DNMT3A, and DNMT3B participate in DNA methylation in mammals to regulate embryo development, cell differentiation, gene transcription, and other normal biological functions. Abnormal functions of DNMTs are associated with tumorigenesis and other diseases.^[7–8]^

DNMTs were the first epigenetic targets for which inhibitors received the approval of the Food and Drug Administration (FDA) of the USA: the nucleoside analogs 5-azacitidine (Vidaza) and decitabine or 5-aza-2’-deoxycytidine (Dacogen) (Figure 1), approved in 2004 and 2006, respectively, for the treatment of the myelodysplastic syndrome.^[9]^ DNMTs are promising epigenetic targets for the treatment of several types of cancer, including acute myeloid leukemia, colorectal, pancreatic, lung, ovarian, and breast cancer, which have been reviewed comprehensively.^[8, 10]^ Furthermore, DNMTs are also attractive targets for the investigation or treatment of other diseases such as diabetes,^[11]^ autoimmune,^[12]^ and neurological disorders.^[13]^ Inhibitors of DNMTs are also emerging as programs to develop combination therapies in drug cocktails or compounds targeting more than one epigenetic target simultaneously.^[14]^ For instance, Rabal et al. reported recently dual inhibitors of DNMT1 and G9a histone methyltransferase.^[15–16]^ Yuan et al. described a dual DNMT and HDAC inhibitor.^[17]^ Hydralazine is an antihypertensive drug and a week inhibitor of DNMT1 (Figure 1). This compound has been proved in several cancer cell lines, and there are researches that demonstrated that is an inhibitor of DNMT1 with an IC_50_ of 2 μM for A549 cell line (human lung cancer cell line harboring wild-type p53), an IC_50_ of 20 μM for U373MG cell line (human glioblastoma cell line harboring inactive mutant p53)^[18]^ and an IC_50_ of 30 μM for Hut78 cell line (cutaneous T-cell lymphoma).^[19–20]^

**Figure 1.**
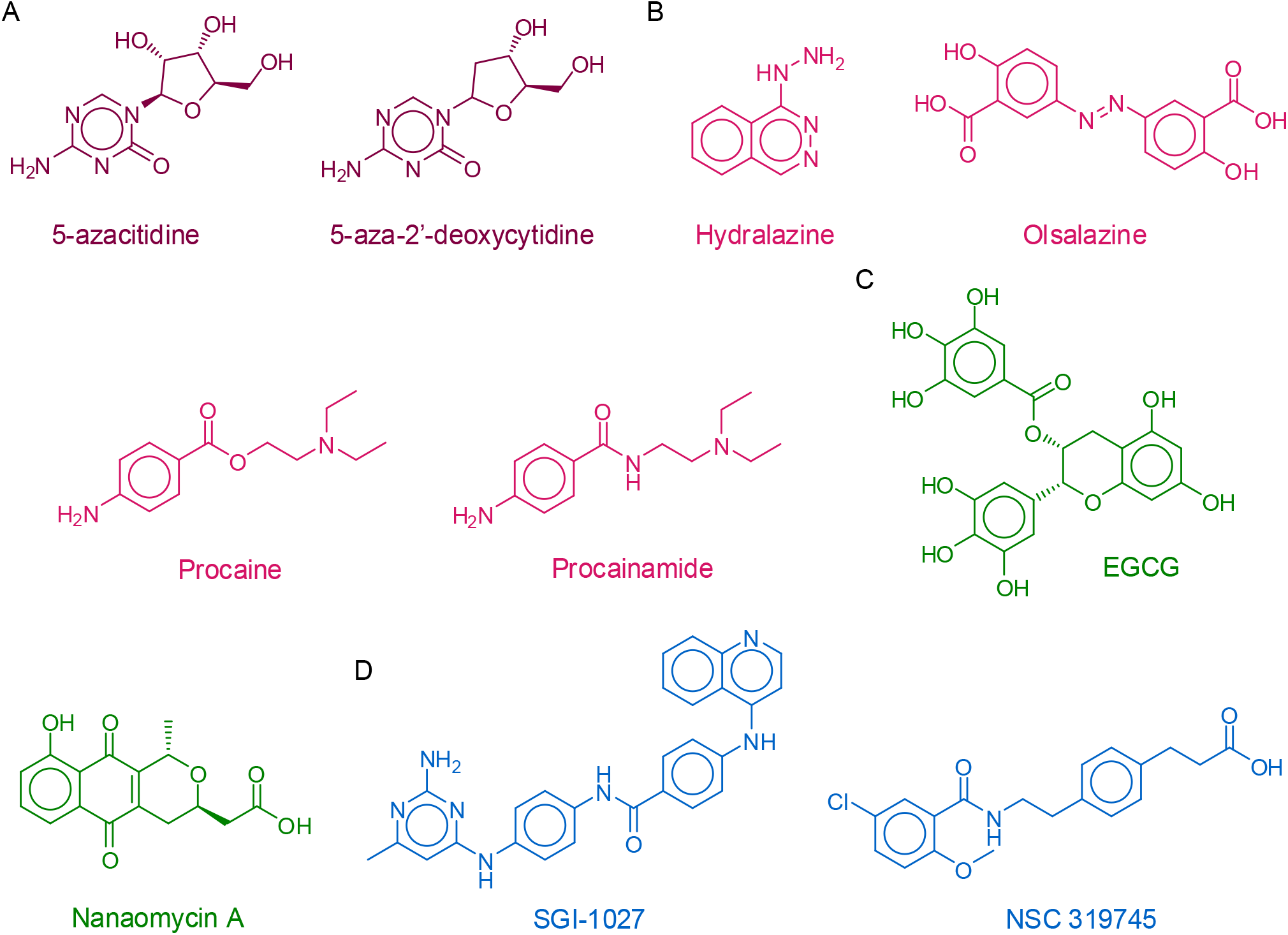
Chemical structures of representative DNMT inhibitors and other compounds with associated hypomethylating properties, from different sources. **A**) approved drugs; **B**) drug repurposing; **C**) Natural products and dietary components; **D**) Small molecules: synthetic compounds.

Despite the fact there are two DNMT inhibitors approved for clinical use, both azacitidine and decitabine have low specificity, poor bioavailability, and instability in physiological conditions and toxicity. Therefore, it has been the interest of our^[21–28]^ and several other research groups^[29–36]^ to identify DNMT inhibitors with novel chemical scaffolds for further development. Inhibition of DNMTs remains a major topic of research not only because of its potential therapeutic benefits but also to understand the essential mechanisms of epigenetic events in cells. There are currently more than 256 compounds in annotated public chemical databases^[37]^ with measured activity vs. DNMTs. Figure S1 in the Supporting Information shows the most frequent scaffolds of active molecules. Figure 1 shows the chemical structures of representative DNMT inhibitors or compounds with DNA demethylation activity from different sources, including drugs for other indications, screening compounds from synthetic origin, and natural products.^[38–40]^ Also, there are several compounds from dietary origin.^[41–43]^ Of note, strong evidence indicates that environmental factors and nutrients play a major role in establishing epigenetic mechanisms, including irregular DNA methylation patterns. Thus, a regular uptake of DNA demethylating agents (which are not necessarily very potent DNMT inhibitors) is hypothesized to have a chemopreventive effect.^[44]^

The DNMT inhibitors have been identified from different approaches or their combination^[44–45]^ such as virtual screening,^[21,30]^ high-throughput screening, lead optimization,^[34,46]^ and structure-guided design^[36]^ to name a few. Amongst the most promising inhibitors are molecules with “long scaffolds” such as the quinolone-based **SGI-1027** (Figure 1) and analogs. With the aid of molecular modeling, it has hypothesized that such compounds with long scaffolds occupy the catalytic site and SAM’s cavity.^[47]^ It has also been proposed that analogs of **SGI-1027** exert their mechanism through interaction with DNA.[48]

As part of an ongoing effort to identify novel DNMT1 inhibitors from different sources and further increase the availability of novel scaffolds, herein, we report five new inhibitors of DNMT1 with distinct chemical scaffolds. Two compounds, glyburide, and panobinostat are approved drugs with potential drug-repurposing applications. In particular, panobinostat is a pan histone deacetylase inhibitor (HDAC), another major epigenetic target, and could be used as a dual epigenetic agent.

## 2. Experimental Section and Computational Methods

We aimed to test compounds from different sources: approved drugs, synthetic compounds from a DNMT focused library, and one compound from dietary origin. We experimentally tested ten molecules with diverse chemical scaffolds and chemical structures different from reported DNMT inhibitors (Figure 2). Thus, we screened in biochemical assays as inhibitors of DNMTs two approved drugs with “long” chemical scaffolds (Figure 2A): the sulfonylurea glybenclamide, approved for the treatment of diabetes (*vide infra*), and panobinostat, a non-selective and potent zinc-dependent HDAC inhibitor (the most potent deacetylase inhibitor on the market), approved in 2018 by the FDA for the treatment of multiple myeloma. Of note, currently epigenetic multitargeting is focused on the inhibition of zinc-dependent HDACs as one of action mechanisms.^[14]^ Due to the known activity of polyphenols and previous evidence of theaflavin activity with DNMT3A,^[49]^ we hypothesized that theaflavin, a dietary component (Figure 2B) present in the back tea, is an inhibitor of DNMT1 and DNMT3B. We also theorized that the naphthalene sulfonamide **7936171** (Figure 2C) with a long scaffold is an inhibitor of DNMT1 and DNMT3B.^[50]^ Finally, we tested six molecules from a DNMT focused library of synthetic molecules (Figure 2D), which are becoming attractive to experimentally screen molecules.^[6]^ To select the ten compounds in this work, commercial availability and reasonable price (the latter an important factor considering the current economic situation imposed by the COVID-19 pandemic) were additional criteria.

**Figure 2.**
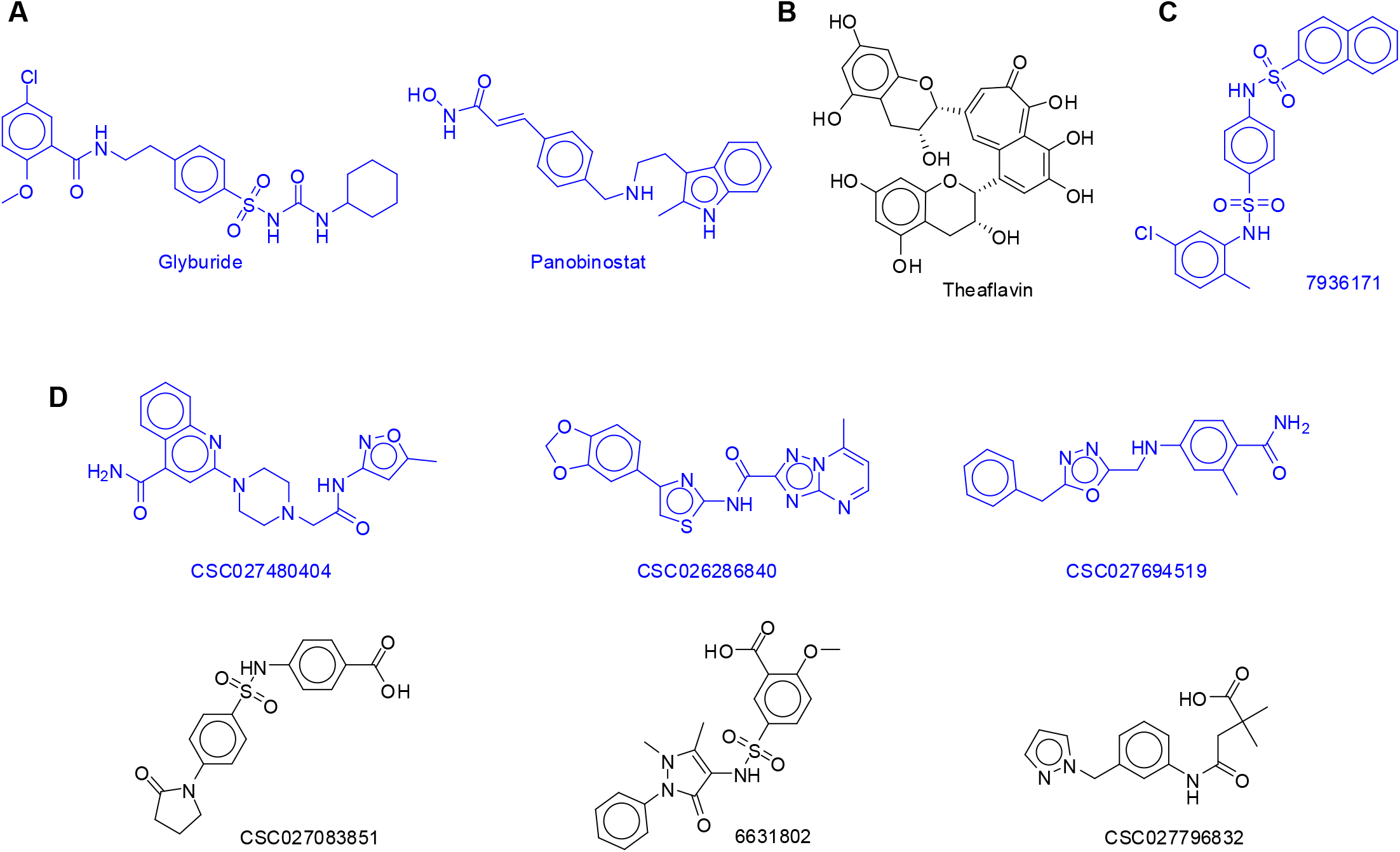
Chemical structures of ten compounds from different sources experimentally tested. Compounds are grouped by their source **A)** approved drugs; **B)** dietary source (natural product); **C)** Inhibitor of the viral NS5 RNA methyltransferase; **D)** DNMT focused library.

First, the ten compounds were tested at a single concentration (100 μM) in duplicate in a biochemical DNMT assay. SAH was used as a reference. The five most active molecules at one-single concentration were further tested in a dose-response manner to obtain the half-maximal inhibitory concentration (IC_50_). Predicted binding modes for the five compounds were studied using molecular docking, implementing a novel re-scoring algorithm recently developed.^[51]^

### 2.1 Compounds

All compounds were purchased from chemical vendors. Glyburide, panobinostat, and theaflavin were purchased from TargetMol. Molecules **7936171** and **6631802** were purchased from Chembridge Corporation. Molecules **CSC027480404**, **CSC026286840**, **CSC027694519**, and **CSC027083851** were acquired from ChemSpace. The compound purity confirmed by the chemical vendors was equal or higher than 90 % (Table S1 in the Supporting information).

### 2.2 Biochemical DNMT assays

The inhibition of the enzymatic activity of DNMT1, DNMT3B, and DNMT3B/3L was tested using the HotSpot^SM^ platform for methyltransferase assays available at Reaction Biology Corporation.^[52]^ HotSpot^SM^ is a low volume radioisotope based assay that uses tritium-labeled AdoMet (^3^H-SAM) as a methyl donor. The test compounds diluted in dimethyl sulfoxide were added using acoustic technology (Echo550, Labcyte) into enzyme/substrate mixture in the nano-liter range. The corresponding reactions were commenced by the addition of ^3^H-SAM, and incubated at 30°C. Total final methylations on the substrate (Poly dI-dC in DNMT1 assay, and Lambda DNA in DNMT3B; DNMT3B/3L assay) were identified by a filter binding method implemented in Reaction Biology. Data analysis was conducted with Graphed Prism software (La Jolla, CA) for curve fits. Reactions were carried out at 1 μM of SAM. In all assays, SAH was used as a standard positive control. The ten compounds were tested first with DNMT1 and DNMT3B at one 100 μM concentration in duplicate. The five most active compounds were tested in 10-dose IC_50_ (effective concentration to inhibit DNMT1 activity by 50%) with a three-fold serial dilution starting at 100 μM. Theaflavin was also tested with DNMT3B and DNMT3B/L in 10-dose IC_50_ with three-fold serial dilution starting at 100 μM. The authors have previously contracted the screening services of Reaction Biology Corporation to identify a novel inhibitor of DNMT1.^[23]^

### 2.3 Molecular docking and re-scoring

The five compounds tested in a dose-response manner, and that showed activity with DNMT1 (*vide infra*) were docked with this enzyme using the program Molecular Operating Environment (MOE), version 2018.08.^[53]^ The chemical structures of the five compounds were built with MOE. The docking was carried out with the crystal structure of the catalytic domain of DNMT1 obtained from the Protein Data Bank^[54]^ PDB ID: 4WXX.^[55]^ This crystal structure is in complex with SAH and has a resolution of 2.62 Å. The structure of the protein was prepared with the “QuickPrep” tool of MOE v.2018 using the parameters established by default, which help to remove the molecules of structural water and add hydrogens atoms to the protein. In this process, the co-crystallized SAH was removed for the binding site to realize a direct docking. The docking was conducted using default parameters in MOE. Before docking the five newly tested compounds, the docking protocol was validated by re-docking the SAH obtaining a root-mean-square deviation of 1.3 Angstroms (and a docking score of −8.96 kcal/mol).

Docking poses obtained with MOE were re-scored with the Extended Connectivity Interactions Features (ECIF) method recently developed. Briefly, ECIF is a recently reported set of descriptors to represent protein-ligand complexes. These descriptors are defined as a set of protein−ligand atom-type pair counts relying on a detailed description of the connectivity of the atoms involved. It has been shown that machine-learning scoring functions built on ECIF and Gradient Boosting Trees, consistently outperform the performance of scoring functions reported so far regarding the obtention of binding scores in a linear correlation with experimental data, particularly when a distance cutoff criterion of 6 Angstroms is used to derive the descriptors, and purely ligand-based descriptors are added (ECIF6::LD-GBT).^[51]^ From the poses obtained from MOE, the best-ranked one for each compound was prepared using X-Tool^[56]^ and standardized using Standardizer, JChem 20.11.0, 2020, ChemAxon to perceive aromaticity in an interpretable way for the model. All poses were re-scored using the ECIF6::LD-GBT model.^[51]^

## 3. Results

### 3.1 Biochemical DNMT assays

Table 1 summarizes the relative enzymatic activities of DNMT1 and DNMT3B in the presence of 100 μM compound. Compounds that had more than 20% inhibition were regarded as inhibitors and were moved forward to dose-response evaluations. We used a similar criterion in a previous identification of novel chemical scaffolds.^[21]^ Seven molecules showed detectable inhibition with DNMT1, of which the five most active were theaflavin (65 % inhibition), **CSC027694519** (65 %), panobinostat (63 %), **7936171** (62 %), glyburide (60 %). The least active were **CSC027480404** (29 %) and **CSC026286840** (27 % inhibition).

**Table 1.**
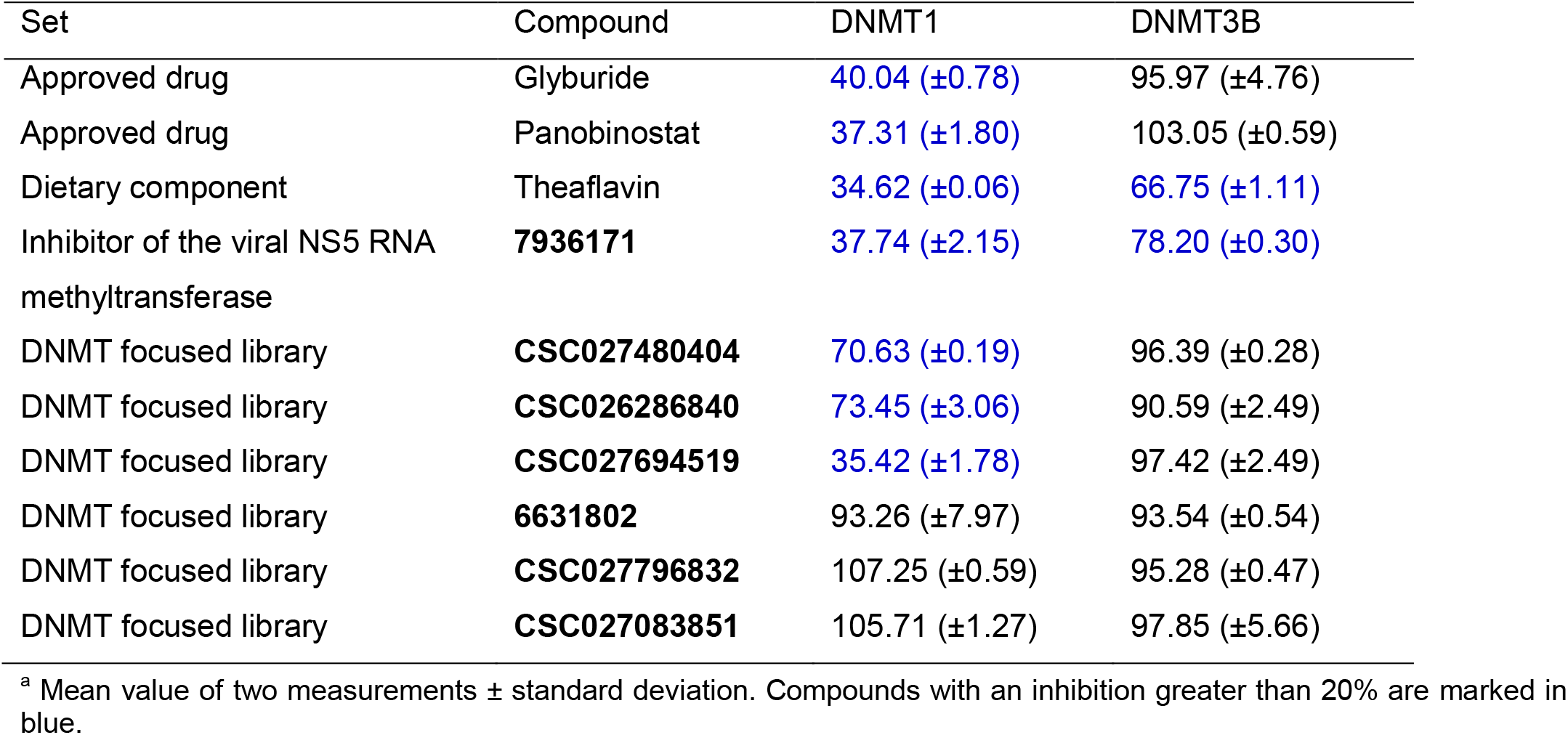
Results of the relative enzymatic activity of DNMT1 and DNMT3B in percent.^a^.

Theaflavin and **7936171**, showed detectable inhibition with DNMT3B (33 and 22 % inhibition, respectively). All other eight molecules were inactive with DNMT3B.

The five compounds with the highest percentage of inhibition at 100 μM with DNMT1 were tested in dose-response manner. Theaflavin that showed the best activity at a single dose with DNMT1 and DNMT3B was also tested drug-response manner with DNMT3B and DNMT3B/3L. Table 2 summarizes the results.

**Table 2.** Results of dose-response evaluations for selected compounds (IC_50_) with DNMT1.^a^.

| Set | Compound | DNMT1 (IC<sub>50</sub> μM) |
| --- | --- | --- |
| Approved drug | Glyburide | 55.85 (±1.11) |
| Approved drug | Panobinostat | 76.78 (±0.23) |
| Dietary component | Theaflavin<sup>b</sup> | 85.33 (±0.14) |
| Inhibitor of the viral NS5 RNA methyltransferase | **7936171** | 78.53 (±8.60) |
| DNMT focused library | **CSC027694519** | 85.11 (±4.10) |

<sup>a</sup> Mean value of two measurements. SAH was included as a positive control (IC<sub>50</sub> of 0.26 μM); <sup>b</sup> Also evaluated with DNMT3B and DNMT3B/3L. The IC<sub>50</sub> was > 100 μM

The two approved drugs, glyburide and panobinostat, inhibited DNMT1 with IC_50_ values of 55.85 and 76.78 μM, respectively. Theaflavin had an IC_50_ value of 85.33 μM. The other two small-molecules **793617** and **CSC027694519** had IC_50_ values of 78.53 and 85.1 μM, respectively. SAH was used as a non-specific positive control and confirmed its effective inhibition of DNMT1 with an IC_50_ of 0.26 μM under the assay’s conditions. Theaflavin, which showed the best activity at a single dose with DNMT1 and DNMT3B (Table 1), was also tested in a dose-response manner with DNMT3B and DNMT3B/3L showing, in both cases, IC_50_ values > 100 μM.

### 3.2 Molecular docking and re-scoring

The docking was done in MOE v.2018 for the 10 compounds (shown in Figure 2) in the active site of the crystal structure of the catalytic domain of DNMT1 (PDB ID: 4WXX), as described in the Methods section. The docking scores ranged from −8.35 to −7.09 kcal/mol. The scores for the five compounds tested in the biochemical assays in dose-response evaluations ranged from −8.20 to −7.42 kcal/mol. Table 3 summarizes of docking with MOE and re-scoring with (Methods section).

**Table 3.**
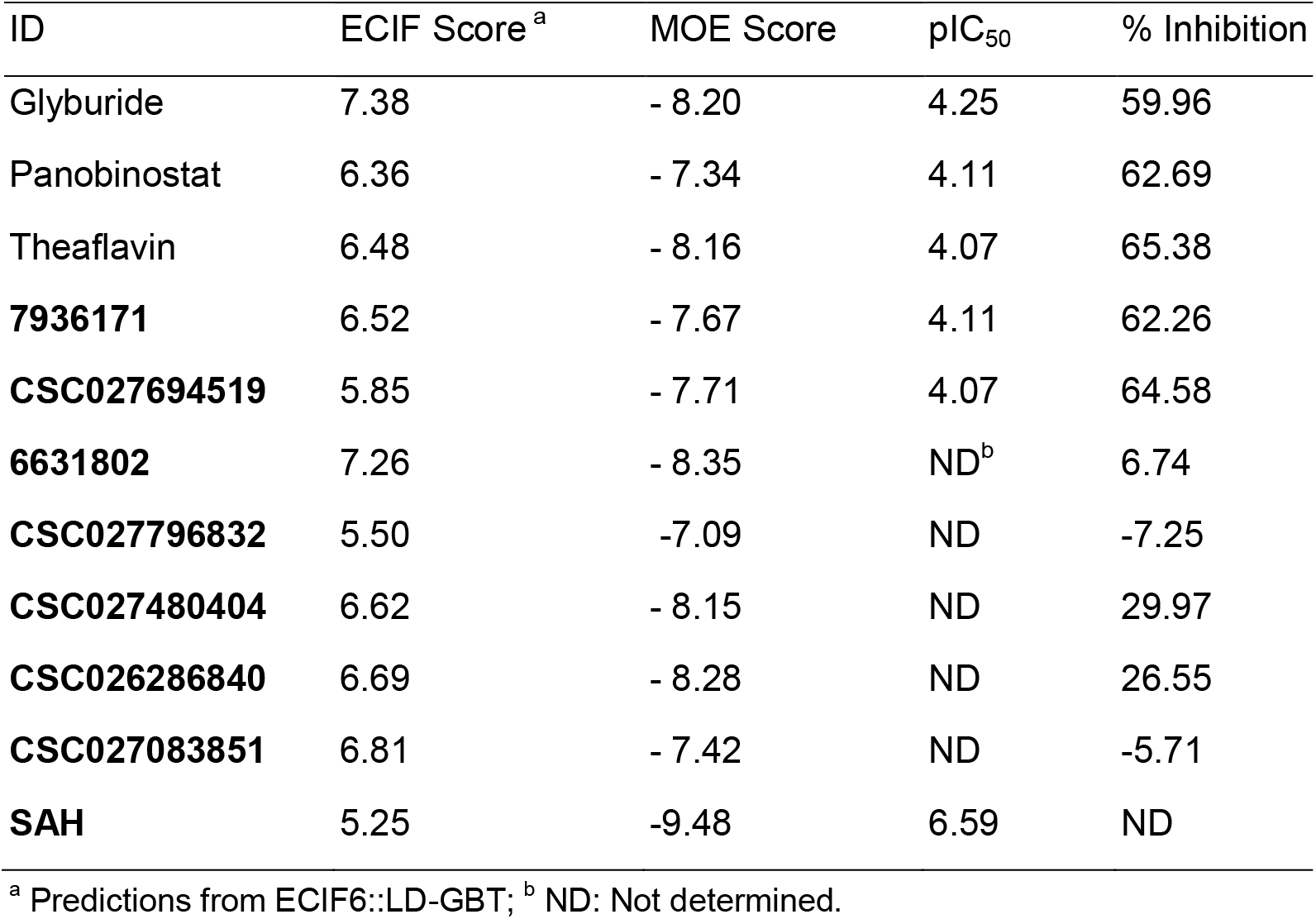
Summary results of docking with Molecular Operating Environment (MOE) and re-scoring with Extended Connectivity Interaction Features (ECIF).

| ID | ECIF Score<sup>a</sup> | MOE Score | pIC<sub>50</sub> | % Inhibition |
| --- | --- | --- | --- | --- |
| Glyburide | 7.38 | - 8.20 | 4.25 | 59.96 |
| Panobinostat | 6.36 | - 7.34 | 4.11 | 62.69 |
| Theaflavin | 6.48 | - 8.16 | 4.07 | 65.38 |
| **7936171** | 6.52 | - 7.67 | 4.11 | 62.26 |
| **CSC027694519** | 5.85 | - 7.71 | 4.07 | 64.58 |
| **6631802** | 7.26 | - 8.35 | ND<sup>b</sup> | 6.74 |
| **CSC027796832** | 5.50 | -7.09 | ND | -7.25 |
| **CSC027480404** | 6.62 | - 8.15 | ND | 29.97 |
| **CSC026286840** | 6.69 | - 8.28 | ND | 26.55 |
| **CSC027083851** | 6.81 | - 7.42 | ND | -5.71 |
| **SAH** | 5.25 | -9.48 | 6.59 | ND |

<sup>a</sup> Predictions from ECIF6::LD-GBT; <sup>b</sup> ND: Not determined.

Figure 3 shows the 2D interaction maps of the predicted binding mode between the human catalytic domain of DNMT1 and the five compounds evaluated in a dose-response manner: glyburide, panobinostat, theaflavin, **793617**, and **CSC027694519.** These interactions correspond to the docking poses with the most favorable docking scores, as calculated with MOE. All the compounds had predicted interactions with catalytic residues and showed hydrogen bond interactions with Glu 1266. Furthermore, glyburide, panobinostat, theaflavin, and **CSC027694519** also showed π-H interactions.

**Figure 3.**
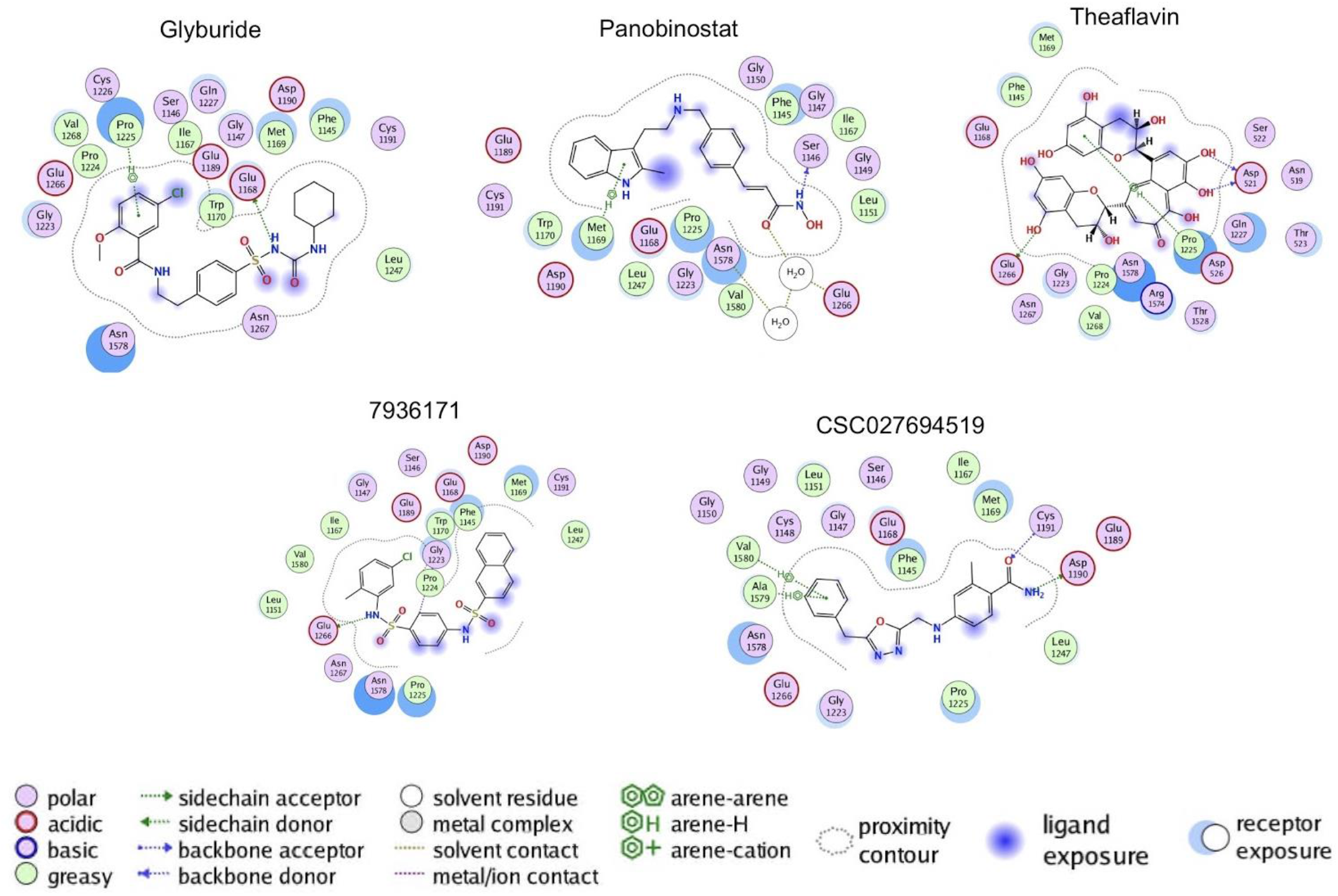
Binding poses predicted with the Molecular with Molecular Operating Environment v.2018 of the five active compounds with the catalytic domain of DNA methyltransferase 1.

The docking poses of the five compounds were re-scored with ECIF, as described in the Methods section. Table 3 summarizes the results of the predictions from ECIF6::LD-GBT for the docking poses generated with MOE. For the five compounds tested in a dose-response manner, the table shows the results of the IC_50_ as the −log value (pIC_50_). The table indicates that among the five compounds with pIC_50_ values, the most active, glyburide, is predicted correctly by the ECIF re-scoring method. Also, the re-scoring of MOE poses ranks glyburide as the one with the highest affinity overall, while MOE score predicts a higher affinity for compounds **6631802** and **CSC026286840**. Figure 4 shows the association between the experimental pIC_50_ of the five compounds evaluated in biochemical inhibition assays of DNMT1 with the MOE’s docking scores (Figure 4A) and re-scoring with ECIF6::LD-GBT (Figure 4B). Clearly, the ECIF re-scoring scheme improved the correlation with the experimental pIC_50_ values.

**Figure 4.**
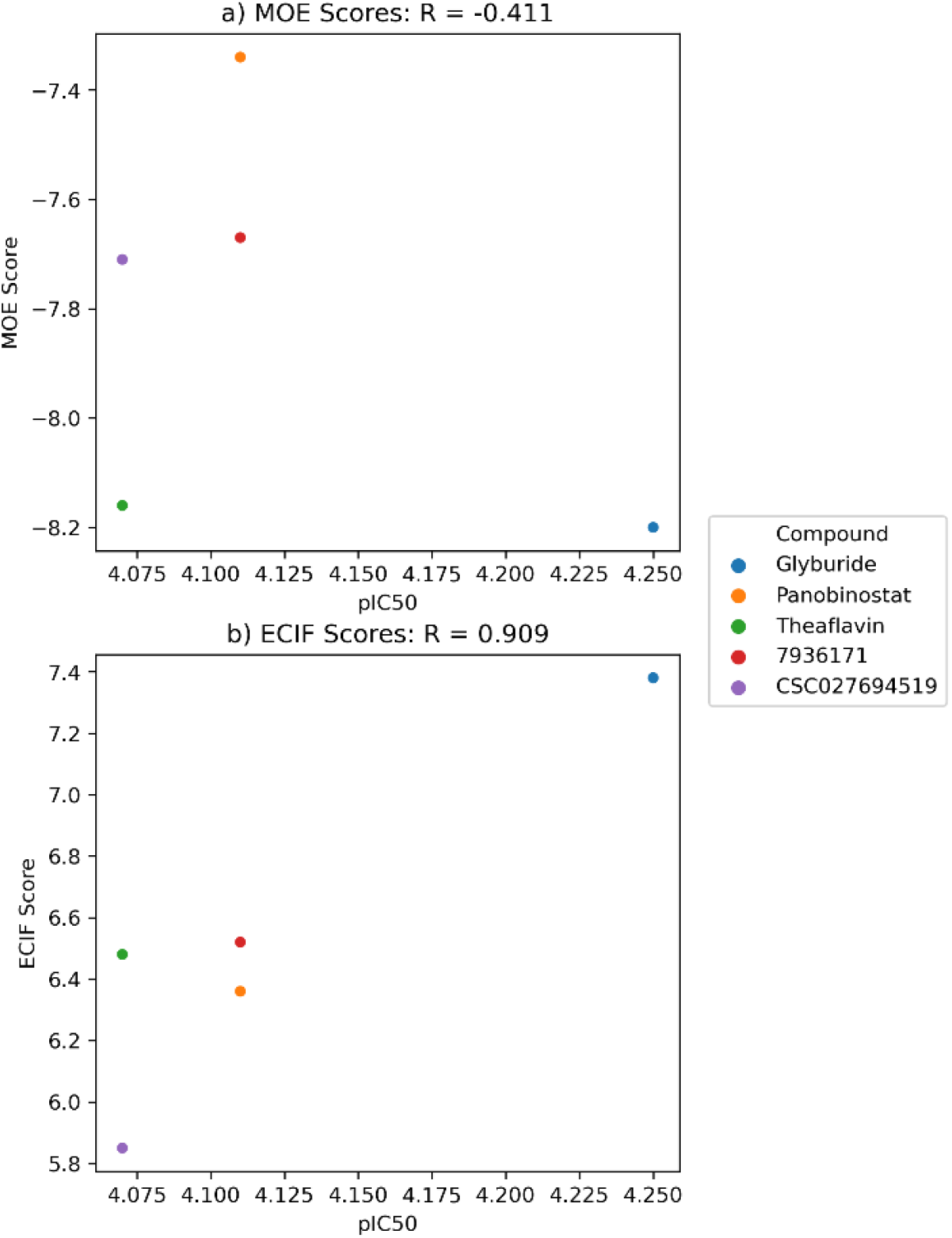
Correlation between the experimental pIC_50_ of the five compounds evaluated in biochemical inhibition assays of DNMT1 with **A)** the docking scores calculated with MOE, and **B)** the ECIF6::LD-GBT scores.

## 4. Discussion

### 4.1 Biochemical DNMT assays

The initial goal of this study was to identify novel small molecule DNMT1 and DNMT3B inhibitors. In this work, we deprioritize testing with DNMT3A because it has been reported that this enzyme can act as both an oncogene and as a tumor suppressor gene, at least in lung cancer (it remains to test if this paradox of DNMT3A applies to other cancers).^[8]^ Seven molecules with detectable inhibition of DNMT1 and two with DNMT3B were initially identified at single dose concentrations (Table 1). An approved drug for the treatment of type 2 diabetes (glyburide) and an approved anticancer drug known as HDAC inhibitor (panobinostat) were among the active compounds with DNMT1. Other active compounds against DNMT1 were a dietary component (theaflavin), a naphthalene sulfonamide (**7936171**) previously reported to be an inhibitor of dengue virus methyltransferase,^[50]^ and three small molecules from a DNMT focused library (**CSC027480404**, **CSC026286840**, and **CSC027694519**).

Overall, the tested compounds showed less activity with DNMT3B (Table 1). Only two compounds, the dietary component theaflavin, and the naphthalene sulfonamide, showed inhibition higher than 20 % with DNMT3B at high concentration. Moreover, theaflavin showed the highest percentage of inhibition (33 %), which was lower than the highest percentage of inhibition showed for DNMT1 (65 %). None of the two compounds had an IC_50_ lower than 100 μM with DNMT3B.

Dose-response assays for the most active compounds at a single dose (Table 2) revealed that, under the assay conditions used in this study, the two approved drugs glyburide and panobinostat had the lowest IC_50_ values (55.85 and 76.78 μM, respectively). The other three active compounds (**7936171**, theaflavin, and **CSC027694519**) had IC_50_ values between 78.5 and 85 μM. Overall, these IC_50_ values are relatively high. However, it should be noted that standard biochemical assays for DNMT enzymes have not been fully established and the results show a large variation between assay conditions. This point has been largely discussed in the literature for DNMT inhibitors.^[47, 57]^ In this work, we used SAH as a positive control, since SAH is a very well-known inhibitor of DNMT.^[58]^ Even though the active compounds’ potency values are not high, all compounds have chemical scaffolds different from the scaffolds reported for known DNMT inhibitors.

Glyburide (Figure 2) is a sulfonylurea. In 1984 it received approval by the United States FDA for the treatment of patients with diabetes mellitus type II. To the best of our knowledge, there are no reports of this compound as an inhibitor of DNMT1. However, several studies support that sulfonylureas are potential anticancer drugs candidates for several cell lines such as colon cancer, human ovarian cancer, kidney cancer, melanoma, and lung cancer.^[59–60]^ Therefore, we propose that glyburide could be further investigated and developed as an epi-drug with potential anticancer activity.

Panobinostat is an orally available pan-deacetylase inhibitor with broad antitumor activity.^[61]^ It has shown inhibitory activity vs. different types of cancer, for example, malignant glioma cells with an IC_50_ range of 0.3 μM to 0.23 μM,^[62]^ carcinoma cells (IC_50_ between 0.4 nM – 1.3 nM),^[63]^ and in human erythroleukemic cell lines (IC_50_ between 40 nM to 51 nM).^[64]^ One of the principal benefits of the panobinostat is that being an HDAC inhibitor will avoid the histone acetylation, which helps to maintain closed the chromatin and decrease the gene transcription involved with proliferation, differentiation, and progress of the cancer cells. Zopf et al. reported that panobinostat reduced DNMT1 (and DNMT3a) activities and expression in liver cancer cell lines.^[65]^ In the study of Zopf et al., it was concluded that inhibitors of HDACs can indirectly control DNA methylation. In this work, it was confirmed that panobinostat also inhibits the enzymatic activity of DNMT1 directly. Of note, there is a recent interest in developing dual inhibitors of DNMTs and HDACs, and results in this direction look promising.^[17]^ Indeed, Yuan et al. reported hydroxamic acid derivatives of a small molecule previously identified as a week inhibitor of DNMT1 from virtual screening of a large chemical library.^[21]^ In the work of Yuan et al., the hydroxamic acid derivatives of NSC 319745 showed inhibition of DNMT1, HDAC1, and HDAC6, plus cytotoxicity activity against human cancer cells.^[46]^ In a second example in the clinic, the combined use of decitabine (a DNMT inhibitor) and belinostat (an HDAC inhibitor) increases the chemotherapy efficiency.^[66]^ Besides, Min et al. showed that an increase in transcription of DNMT1 is one of the mechanisms of resistance of anti-cancer drugs targeting HDACs, such as vorinostat. Consequently, it has been shown that co-targeting DNMT1 improved the antitumor efficacy of vorinostat and other HDAC inhibitors.^[67]^ These contributions, including the findings of this work, are in line with the epigenetic multitargeting. The advantages of this approach have been demonstrated in both *in vitro* and *in vivo* disease models by the co-administration of an epigenetic agent with another drug.^[68]^

The naphthalene sulfonamide **7936171** (also with ID ZINC 01078518) is a known inhibitor of the viral NS5 RNA methyltransferase with an IC_50_ of 64.2 μM and an EC_50_ of 12 μM.^[50]^ The viral RNA domain involves the use of SAM as the methyl donor and generates SAH as the final product. Compound **7936171** has a “long” and different scaffold from the known DNMT1 inhibitors published so far (Table S1).

Theaflavin is a natural product polyphenol found in green and black tea and coffee (*vide supra*) with previously measured enzymatic inhibitory activity of DNMT3A.^[49]^ However, there were no reports on its inhibitory potential of DNMT1 and DNMT3B. In this work, it was measured for the first time its activity with both enzymes. Despite the fact its IC_50_ is high (85.3 μM, under the assay conditions of this work), this dietary component could contribute to the modulation of DNMT1. Interestingly, it has been proposed that the modulation of normal levels of DNMT could be conveniently achieved through the dietary uptake of food chemicals (or other “safe” natural products). A prominent example of this hypothesis has been suggested for the polyphenol compound from green tea, EGCG (Figure 1), which has been proposed to inhibit DNMT1 and reactivate methylation-silenced genes in cancer.^[44]^

The other two small molecules, **7936171** and **CSC027694519**, had no previous reports of inhibition of DNMTs. As discussed in the next section, both molecules have dug-like characteristics and have novel scaffolds that can be further optimized for DNMT1 inhibition.

### 4.2 Computational studies

Herein we employed *in silico* methods to rationalize at the molecular level the experimental results of the most active molecules. Of the different mechanisms described to inactivate DNMT activity^[69]^ we hypothesized that the herein identified inhibitors are SAM competitors. This hypothesis is based on the “long” scaffolds of the active compounds such as glyburide, panobinostat, **7936171**, and **CSC027694519**. The predicted binding modes suggested strategies for the structure-based optimization of the small molecules. For instance, comparing the predicted binding mode of **CSC027694519** with the co-crystallized position of SAH in PDB ID: 4WXX (Figure S2 in the Supporting Information) suggests that substitution with a polar group in the benzyl ring of **CSC027694519** could enhance the affinity, making polar or hydrogen bond contacts with amino acid residues such as Ser1146, Gly1150, Leu1151, Val1580 (by comparison with the primary amine or carboxylate groups of SAH). Of note, **CSC027694519** already makes hydrogen bond interactions with the polar side chains of Asp1190 and Cys1191 (similar to the hydrogen bond contacts of SAH, Figure S2). Similarly, comparing the predicted binding mode of **7936171** with the co-crystallized position of SAH (Figure S2) suggest that the naphthalene ring of **7936171** could be replaced with polar heteroaromatic rings. Of course, the synthesis and testing of the analogs should be made, which is one of the major perspectives of this work. It remains to confirm the hypothesis of the putative binding site of these molecules that can be further tested experimentally with binding completion assays once more potent compounds are identified.

## 5. Conclusion

DNMTs are a fundamental class of epigenetic regulatory enzymes. In this work, we tested ten compounds with novel scaffolds as inhibitors of DNMT1 and DNMT3B in biochemical assays. Seven compounds showed at least 20% inhibition of DNMT1 at 100 μM. Five molecules showed activity in dose-response inhibition assays with DNMT1 with IC_50_ between 55.8 and 85.3 μM. Although the molecules’ overall potency is not high under this work’s assay conditions, four compounds have novel chemical scaffolds, not previously described as inhibitors of DNMT1 in biochemical assays. Notably, glyburide, an approved drug for diabetes type II treatment, showed the best potency of the compounds tested in this study and can be further investigated in its role in epigenetic mechanisms. Also, glyburide could be pursued for drug repurposing applications. It was also concluded that panobinostat is an inhibitor of the enzymatic activity of DNMT1. Remarkably, panobinostat, an epigenetic drug for treating different types of cancer, has shown inhibition of DNMT1 in liver cancer cell lines.^[65]^ Therefore, it is concluded that panobinostat, a known HDAC inhibitor, can be further investigated as a multitarget epigenetic agent. In this work, we also concluded that theaflavin, a natural dietary product, inhibits DNMT1 and does not show significant inhibition of DNMT3B. Therefore, this work also complements theaflavin’s DNMT activity profile that had reported activity with DNMT3A.^[49]^ It was also concluded that compounds **7936171** and **CSC027694519** and the two compounds that showed 30 % inhibition of DNMT1 at 100 μM (**CSC027480404** and **CSC026286840**), can be starting points of optimization programs to improve their potency.

Molecular docking of the most active compounds helped propose binding modes with the catalytic binding site of DNM1. Although MOE scores predicted SAH as the most potent inhibitor, the novel ECIF re-scoring scheme ECIF6::LD-GBT improved significantly the correlation of the docking scores calculated with MOE with the experimental pIC_50_ values considering the rest of the compounds. Therefore, docking with MOE for compound selection and re-scoring with ECIF for prioritizing experimental tests can guide the optimization programs of the DNMT1 inhibitors identified in this work, which is one of the main perspectives of this work. Another perspective is testing the DNA demethylation activity of the most active compounds. This can be achieved directly with **7936171** and **CSC027694519** or after the compounds are optimized.

## Abbreviations

^3^H-SAM: Tritium-labeled AdoMet
DNMT: DNA methyltransferase
ECIF: Extended Connectivity Interactions Features
FDA: Food and Drug Administration
^3^H-SAM: tritium-labeled AdoMet
HDAC: Histone deacetylase inhibitor
IC_50_: Half maximal inhibitory concentration
MOE: Molecular Operating Environment
PDB: Protein Data Bank
SAH: *S*-adenosyl-*L*-homocysteine
SAM: *S*-adenosyl-L-methionine.

## Supporting information

Chemical structures of the ten most frequent (Bemis-Murcko) scaffolds of the active DNMT inhibitors available in ChEMBL 27 (**Figure S1**); Chemical vendors of the ten compounds tested and their purity as supplied by the vendor (**Table S1**); Comparison of the co-crystal position of SAH in the catalytic site of DNMT1 (PDB ID: 4WXX) with the predicted binding mode of A) CSC027694519 and B) 7936171 (**Figure S2**).

## Acknowledgements

This work was funded by the *Consejo Nacional de Ciencia y Tecnología* (CONACyT), Mexico (grant 282785). KEJ-M thanks the support from CONACyT. NS-C and FDP-M and are also thankful to CONACyT for the PhD scholarships number, 335997 and 660465/576637, respectively. Valuable discussions with Oscar Méndez-Lucio, Jennifer Tamayo, Carlos Velazquez, and Miguel Herrera are highly appreciated.

## Table of Contents

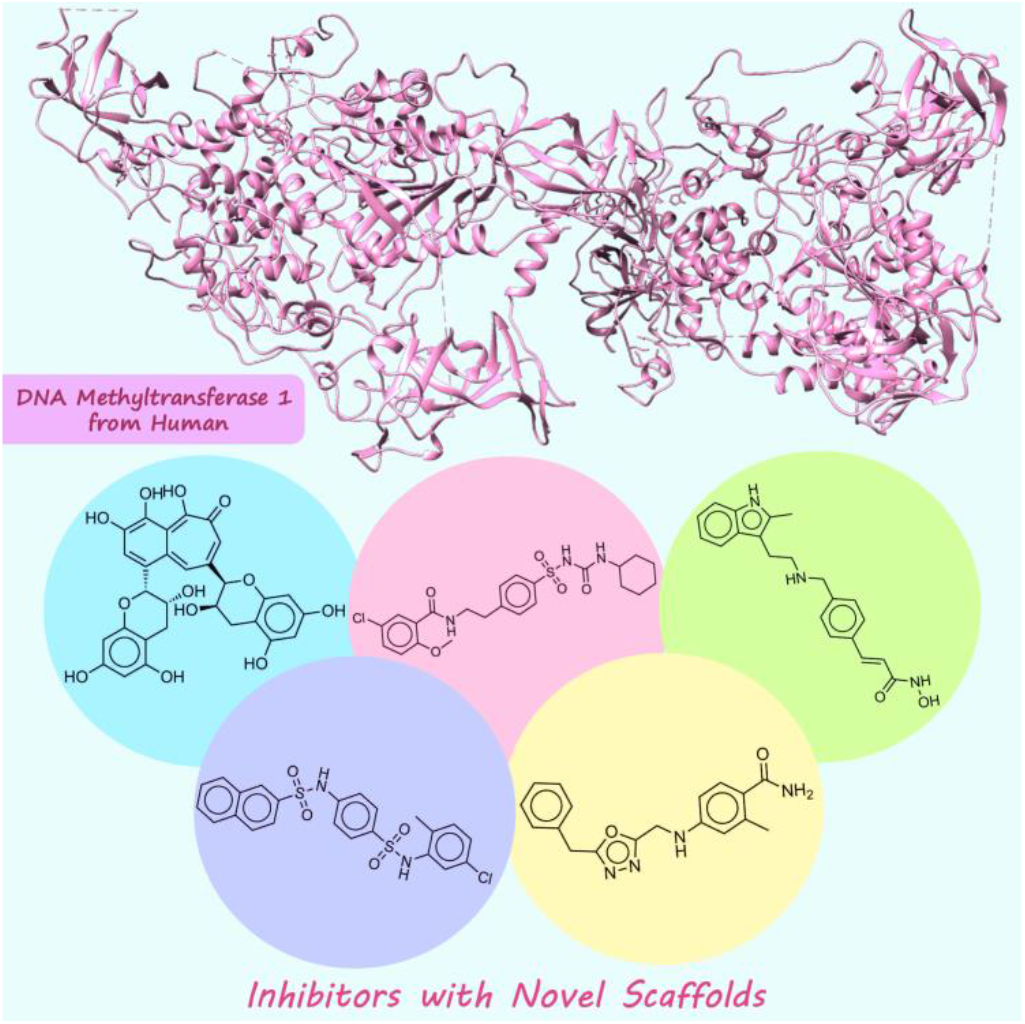

We report five distinct molecules as enzyme inhibitors of DNMT1. One inhibitor was the antidiabetic drug glyburide that could be pursued for drug repurposing. Another inhibitor was the HDAC inhibitor and approved panobinostat and other small molecules with novel scaffolds that can be further optimized.

## Notes

### Competing Interest Statement

The authors have declared no competing interest.

