## Supporting Information for "DNA Methyltransferase Inhibitors with Novel Chemical Scaffolds"

### Contents

| | | Page |
|---|---|---|
| **Figure S1** | Chemical structures of the ten most frequent (Bemis-Murcko) scaffolds of the active DNMT inhibitors available in ChEMBL 27. | S2 |
| **Table S1** | Chemical vendors of the ten compounds tested and their purity as supplied by the vendor. | S3 |
| **Figure S2** | Comparison of the co-crystal position of SAH in the catalytic site of DNMT1 (PDB ID: 4WXX) with the predicted binding mode of **A**) **CSC027694519** and **B**) **7936171**. | S4 |

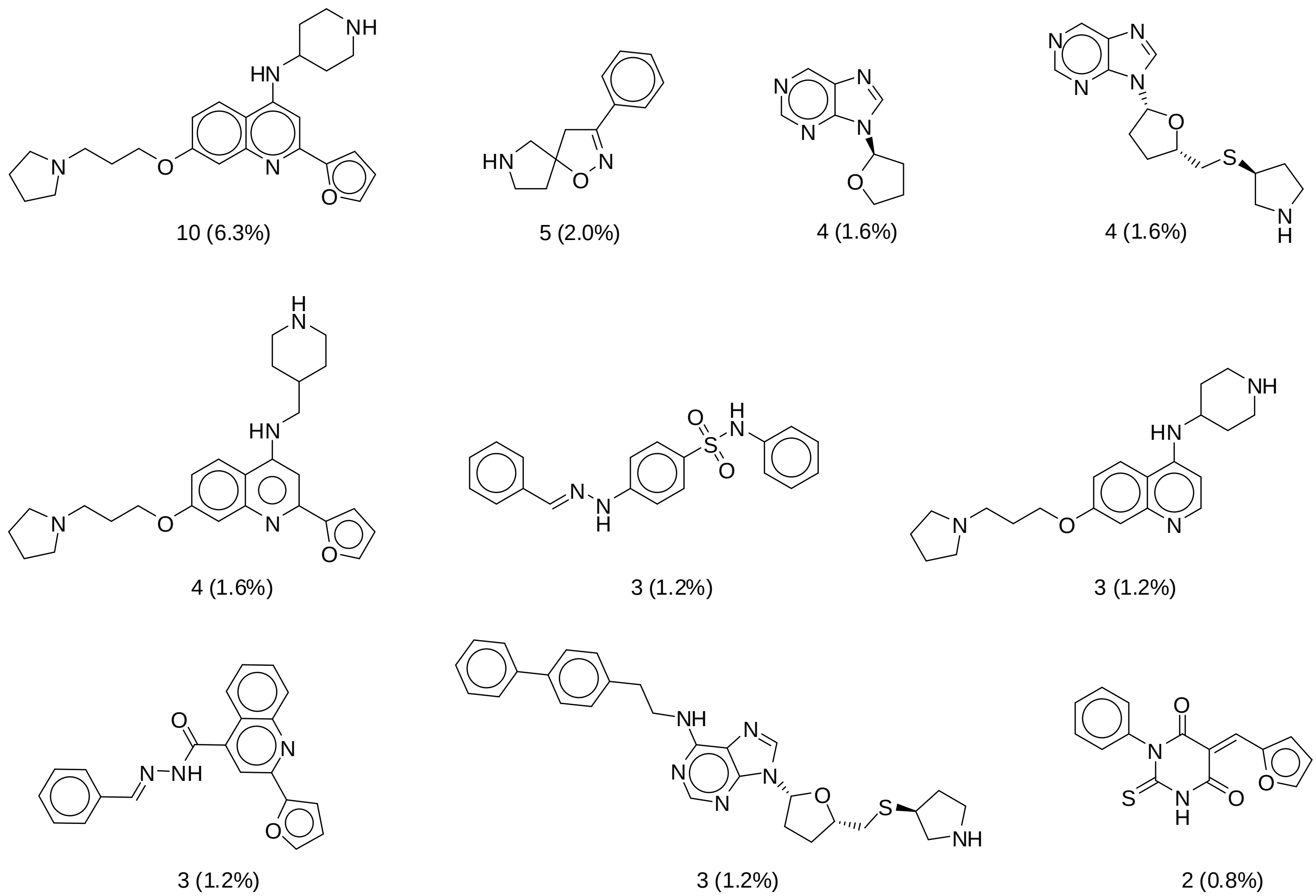


**Figure S1**. Chemical structures of the ten most frequent (Bemis-Murcko) scaffolds of the active DNMT inhibitors available in ChEMBL 27. The percentage of frequency is indicated in parenthesis.

**Table S1.** Chemical vendors of the ten compounds tested and their purity as supplied by the vendor.

| Compound | Vendor | Purity % (provided by vendor) |
|---|---|---|
| Glyburide | TargetMol | 99.77 |
| Panobinostat | TargetMol | 98 |
| Theaflavin | TargetMol | 97.76 |
| **7936171** | Chembridge | >=90 |
| **CSC027480404** | ChemSpace | 90 |
| **CSC026286840** | ChemSpace | 100 |
| **CSC027694519** | ChemSpace | 100 |
| **6631802** | Chembridge | >=90 |
| **CSC027796832** | ChemSpace | 94 |
| **CSC027083851** | ChemSpace | 100 |

**A**

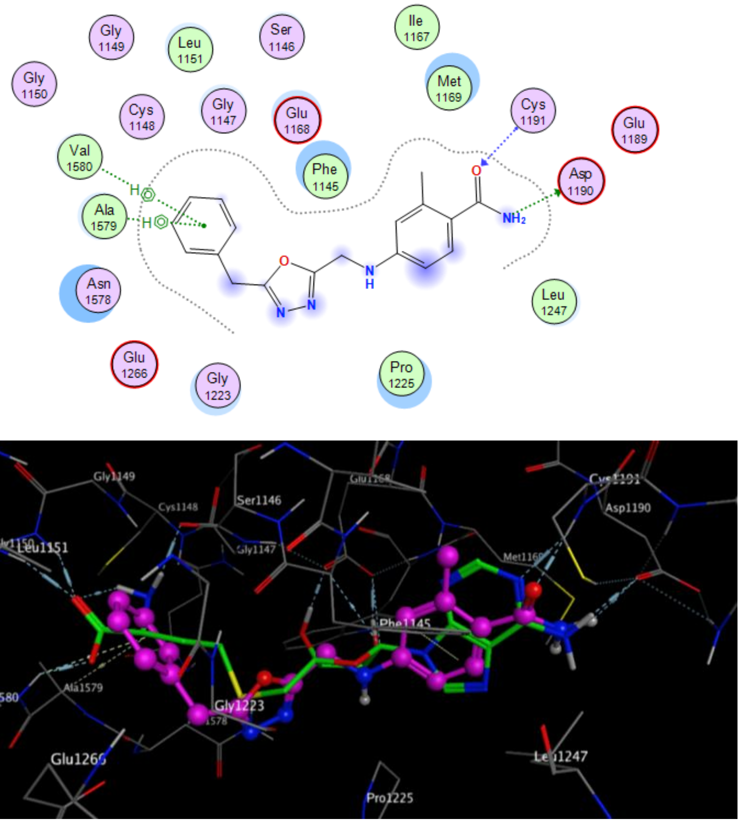


**B**

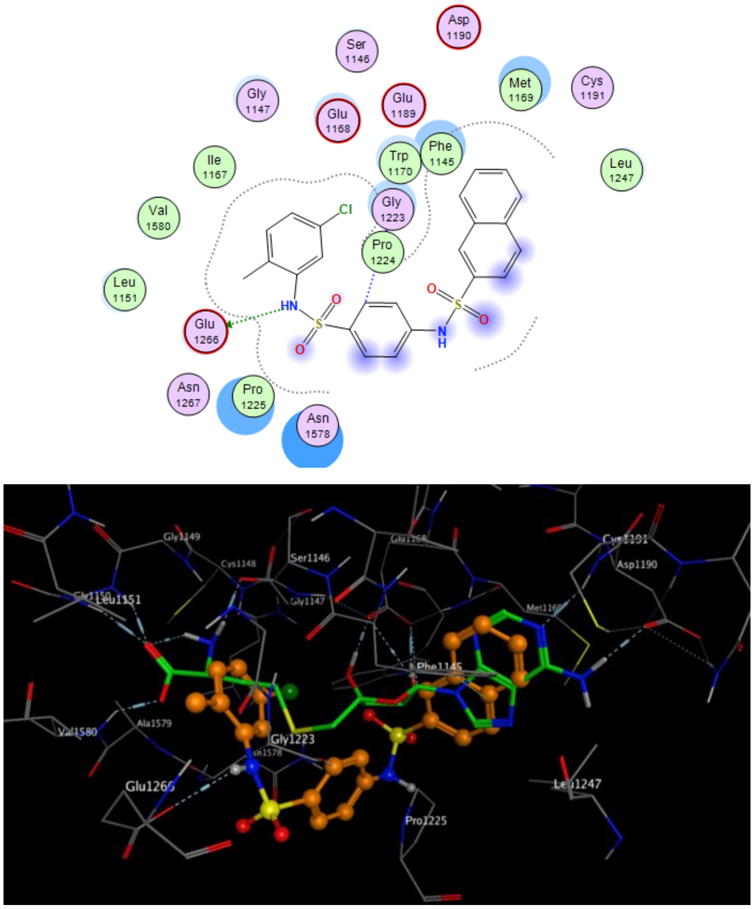


**Figure S2**. Comparison of the co-crystal position of SAH in the catalytic site of DNMT1 (PDB ID: 4WXX) with the predicted binding mode of **A**) **CSC027694519** and **B**) **7936171**.
